# CellPolaris: Decoding Cell Fate through Generalization Transfer Learning of Gene Regulatory Networks

**DOI:** 10.1101/2023.09.25.559244

**Authors:** Guihai Feng, Xin Qin, Jiahao Zhang, Wuliang Huang, Yiyang Zhang, Wentao Cui, Shirui Li, Yao Chen, Wenhao Liu, Yao Tian, Yana Liu, Jingxi Dong, Ping Xu, Zhenpeng Man, Guole Liu, Zhongming Liang, Xinlong Jiang, Xiaodong Yang, Pengfei Wang, The X-Compass Project Consortium, Ge Yang, Hongmei Wang, Xuezhi Wang, Ming-Han Tong, Yuanchun Zhou, Shihua Zhang, Yiqiang Chen, Yong Wang, Xin Li

**Affiliations:** State Key Laboratory of Stem Cell and Reproductive Biology, Institute of Zoology, Chinese Academy of Sciences, Beijing,100101, China; Institute for Stem Cell and Regenerative Medicine, Chinese Academy of Sciences, Beijing, 100101, China; Beijing Institute for Stem Cell and Regenerative Medicine, Beijing, 100101, China; Beijing Key Laboratory of Mobile Computing and Pervasive Device, Institute of Computing Technology, Chinese Academy of Sciences, Beijing, 100190, China; CEMS, NCMIS, HCMS, MDIS, RCSDS, Academy of Mathematics and Systems Science, Chinese Academy of Sciences, Beijing, 100190, China; Computer Network Information Center, Chinese Academy of Sciences, Beijing,100083, China; State Key Laboratory of Multimodal Artificial Intelligence Systems, Institute of Automation, Chinese Academy of Sciences, Beijing, 100190, China; State Key Laboratory of Molecular Biology, Shanghai Key Laboratory of Molecular Andrology, Shanghai Institute of Biochemistry and Cell Biology, Center for Excellence in Molecular Cell Science, University of Chinese Academy of Sciences, Chinese Academy of Sciences, Shanghai, 200031, China; University of Chinese Academy of Sciences, Beijing, 100864, China; School of Software, Yunnan University, Kunming 650091, China

**Author notes:** Correspondence (X. Li), (Y. Wang), (Y. Chen), (S. Zhang), (Y. Zhou), (M. Tong), (G. Feng). These authors contributed equally to this work (Guihai Feng, Xin Qin, Jiahao Zhang, Wuliang Huang, Yiyang Zhang, Wentao Cui). A list of affiliations appears at the end of the paper (The X-Compass Project Consortium).

## Abstract

Cell fate changes are determined by gene regulatory network (GRN), a sophisticated system regulating gene expression in precise spatial and temporal patterns. However, existing methods for reconstructing GRNs suffer from inherent limitations, leading to compromised accuracy and application generalizability. In this study, we introduce CellPolaris, a computational system that leverages transfer learning algorithms to generate high-quality, cell-type-specific GRNs. Diverging from conventional GRN inference models, which heavily rely on integrating epigenomic data with transcriptomic information or adopt causal strategies through gene co-expression networks, CellPolaris employs high-confidence GRN sources for model training, relying exclusively on transcriptomic data to generate previously unknown cell-type-specific GRNs. Applications of CellPolaris demonstrate remarkable efficacy in predicting master regulatory factors and simulating in-silico perturbations of transcription factors during cell fate transition, attaining state-of-the-art performance in accurately predicting candidate key factors and outcomes in cell reprogramming and spermatogenesis with validated datasets. It is worth noting that, with a transfer learning framework, CellPolaris can perform GRN based predictions in all cell types even across species. Together, CellPolaris represents a significant advancement in deciphering the mechanisms of cell fate regulation, thereby enhancing the precision and efficiency of cell fate manipulation at high resolution.

## Introduction

The orchestration of an organism’s development, from a fertilized egg to adulthood, is governed by a sophisticated network of precise genetic programs(de Laat and Duboule, 2013; Peter and Davidson, 2011; Thiery and Sleeman, 2006). These programs collectively form an expansive regulatory network that integrates external signals from the environment and neighboring cells to execute intricate developmental processes (Perrimon et al., 2012). Transcription factors (TFs) serve as core nodes in this network, precisely regulating cell fate by binding to specific elements on target genes, such as promoters or enhancers, in a spatiotemporal-specific manner. TFs play a crucial role in cellular reprogramming and differentiation (Kim and Shendure, 2019; Lambert et al., 2018). Therefore, deciphering TF-centered gene regulatory networks is essential for understanding and manipulating cell fate.

However, TF regulation is generally cell type and stage-specific, making it challenging to determine target genes even through methods such as Chromatin Immunoprecipitation (ChIP) (Park, 2009). With the advent of high-throughput sequencing technologies, researchers have attempted to infer TF-regulated target genes by constructing co-expression networks using scRNA-Seq or bulk RNA-Seq data. However, this method introduces many false-positive connections by approximating correlations as causal relationships (Aibar et al., 2017; Chan et al., 2017; Huynh-Thu et al., 2010). Subsequently, the emergence of ATAC-Seq, which identify open regions including numerous transcription factor binding sites, greatly improve the accuracy of TF binding site and regulatory network predictions. Tools such as PECA2 and its multi-modal version scREG, as well as other tools such as CellOracle and DeepTFni, have significantly improved the accuracy of TF-centered gene regulatory network prediction (Duren et al., 2022; Duren et al., 2020; Kamimoto et al., 2023; Li et al., 2022). However, these tools also require additional information such as input from ATAC-Seq, which limits their applicability.

Previous studies have typically focused on inferring the GRN of a single tissue (Liu et al., 2014). Due to differences in gene expression between tissues, previous deep learning models cannot effectively learn tissue-specific regulatory mechanisms (Mignone et al., 2020; Pio et al., 2022; Yuan and Bar-Joseph, 2019). Generalization algorithms in deep learning can enable the model to fit different tissues adaptively, thereby addressing this issue.

In this study, we present CellPolaris, a deep learning model designed to infer gene regulatory networks (GRNs) solely from RNA-Seq data. Compared to existing tools, CellPolaris uniquely operates without the need for additional data beyond transcriptome information. Instead, it harnesses the power of transfer learning and generalization techniques to integrate pre-existing transcription factor regulatory networks and generate cell type-specific GRNs with precise regulatory weights. Notably, the GRN network generation supports cross-species transfer, making the model applicable for predicting transcription factor perturbations in species lacking regulatory annotations. We emphasize that our approach, as a deep learning model, has a higher potential for performance improvement compared to existing methods as it can utilize high-confidence network inferences generated by other tools for model training.

We apply our transfer GRN method to predict cell type-specific GRNs in cellular reprogramming or differentiation experiments. We then compare the differential networks to screen core TFs for cell fate transformation. The results show that the screened TFs have been applied to existing reprogramming systems. Furthermore, we use the transfer network to analyze single-cell transcriptomic data in the development or differentiation process and develop a Probabilistic Graphical Model (PGM) to predict the impact of TF perturbations on cell fate. We apply the model to the round spermatid differentiation and find that our model accurately predicts the impact of TFs on the developmental processes, with comparable performance to state-of-the-art models and better performance in certain TF predictions.

## Results

### Design and organization of CellPolaris

Our primary objective is to establish cell type-specific gene regulatory networks (GRNs) to decode cell fate control factors. To achieve this goal, we proposed CellPolaris, which consists of two parts: A generalized transfer learning model to establish cell type-specific GRNs, and downstream tasks that utilize these GRNs to identify cell fate regulation factors (Fig 1). First, we constructed high-confidence GRN database centered on transcription factors using matched RNA-Seq and ATAC-Seq data. We then built a generalized transfer learning model by combining the generated GRNs with corresponding transcriptome data from different cell states. This model supports transfer across tissue types, developmental stages, and species, and can establish cell-specific GRNs by simply inputting bulk or single-cell RNA-Seq expression matrices (Fig 1A). Next, we utilized cell type-specific GRNs to perform different downstream tasks related to cell fate regulation factors. 1) Extracting differential GRNs between cell types, we established a model for predicting core factors involved in cell fate transitions (Fig 1B). 2) We built a probabilistic graphical model for predicting transcription factor perturbations during cell differentiation or reprogramming processes based on GRNs with regulatory weights, which can predict the impact of transcription factor perturbations on cell fate and simulating the impact of transcription factor perturbation on the expression levels of both direct downstream target genes and more distant ones (Fig 1C).

**Figure 1.**
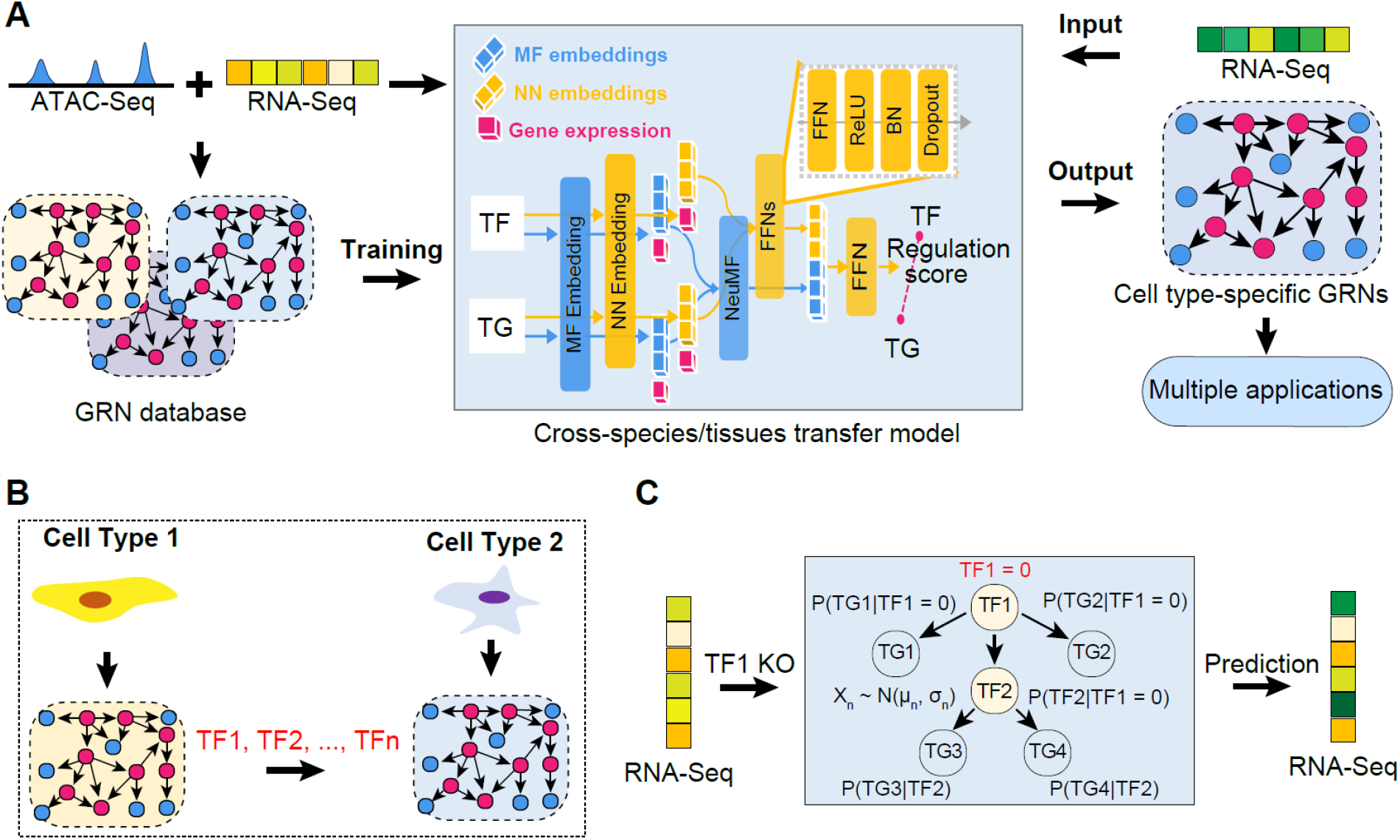
The schematic overview of CellPolaris design. **(A)** Generation of a generalized transfer model using PECA2 tool to construct a GRN database from ATAC-Seq and corresponding cell state RNA-Seq data. This model enables cross-species and cross-tissue analysis. The trained model can take RNA-Seq data as input and generate corresponding GRNs, which can be utilized for downstream applications. **(B)** Prediction of cell fate regulatory factors based on GRN analysis: Comparison of GRN differences between source and target cells during cell fate transitions. Transcription factor nodes in the differential networks were scored and ranked. **(C)** Simulation of the impact of gene perturbation on cell differentiation pathways based on GRN analysis: A probability graphical model of gene expression regulation was constructed using GRN analysis. Simulated knockout genes were set to zero, and the expression changes of neighboring node genes were inferred to deduce changes in cell states.

### GRN construction with transfer learning

The major challenges of cell type-specific GRNs construction are two folds: large data collection cost and out-of-distribution shift. First, constructing cell type-specific GRNs with multi-modal data such as ATAC-seq data and transcriptomic data has considerable cost since these data are collected from complex and repeated in-lab experiments. To solve this problem, we aimed to develop a model that can directly infer cell-type specific GRNs solely from transcriptomic data without relying on matched epigenetic information (Fig 2A). We used high-confidence GRNs as the basis for our model training, and we selected the PECA2 method (Duren et al., 2022; Duren et al., 2020), which was previously developed by our team members, to construct these high-confidence GRNs (Fig S1). This method makes full usage of the transcription factor binding genomic information from ATAC-seq data, the chromatin accessibility activity of regulatory elements, and the gene expression level information from RNA-seq data. Our approach has a distinct advantage: training on limited cellular context with paired expression and chromatin accessibility data, we can get a highly generalized model with the ability to directly constructing GRNs on new context given only transcriptomic data.

**Figure 2.**
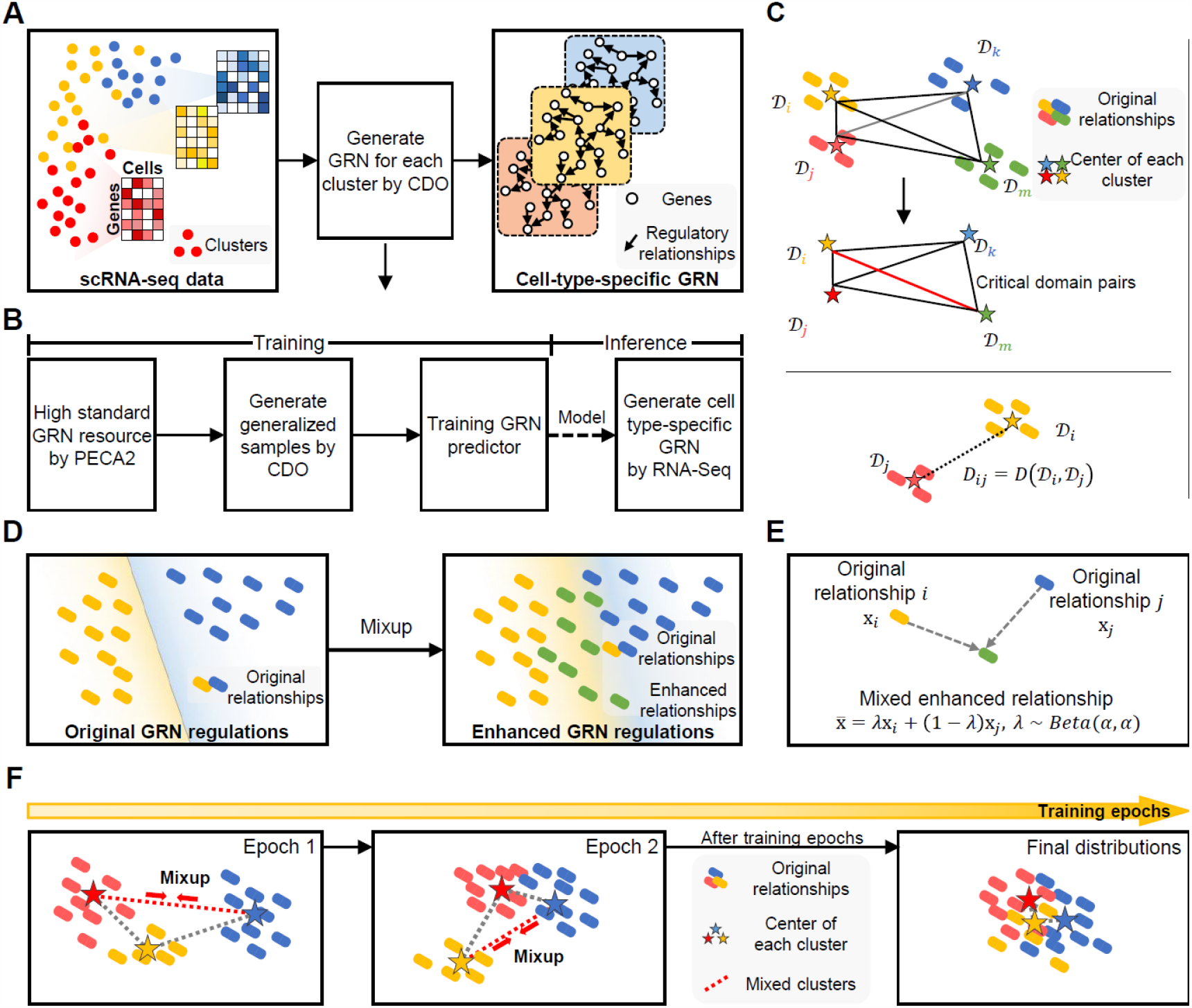
Critical Domain Optimization method. **(A)** The overview of CDO: using the scRNA-seq data as input to generate GRN for each cluster by CDO and output cell-type-specific GRN. **(B)** Workflow of the training and inference of CDO method. Training Phase: Obtain a high-standard GRN resource using PECA2. And then generate generalized samples with CDO for improved training. Finally, train the GRN predictor using the combined data. Inference Phase: Apply the trained model to predict cell type-specific GRNs using RNA-seq data from the target domain. **(C)** Calculate domain similarity with Maximum Mean Discrepancy to dynamically choose the top-σ ratio critical domain pairs, and use mixup to generate generalized samples to emphasize optimizing them. **(D)** Generating Generalized Samples between Critical Domains using Mixup. The left side displays the original relationships, whereas the right side showcases a combination of mixed samples and the original relationships. **(E)** Technical illustration of Mixup between gene relationships from two domains. **(F)** The distribution before (large domain discrepancy exists between domains) and after training with CDO (domain discrepancy is reduced).

Second, the out-of-distribution shift challenge. The GRN regulations in different species, tissues, development periods etc. are dynamic, resulting in data distribution shifts. Most of the existing GRN construction methods usually assume the distribution of the training and testing set are independent and equally distributed (I.I.D). Directly apply the model trained on existing data to new data may get great performance degradation. One widely used and effective approach for out-of-distribution generalization is Mixup (Xu et al., 2020). It performs linear interpolation on sample pairs from two different domains, thereby generating new samples that integrate knowledge from both domains and reducing the distributional gap. However, this method mixes domains randomly, disregarding the complex inherent biological similarity relationships. For example, the similarity between zebrafish and clownfish is higher than that between zebrafish and human. Randomly mixup, on one hand, neglects biologically relevant knowledge critical for learning a unified generalized model, and on the other hand, has higher computational complexity. Motivated by such limitations, we attempted to establish a transfer learning model that can generalize to unseen new transcriptomic data for GRN construction using a critical domain optimization (CDO) strategy (Fig 2B). The core concept of this strategy is to dynamically bridge the expression relationships among the least similar critical domains. The existence of large distribution gaps between these domain pairs makes it notably challenging to acquire invariant knowledge across such diverse domains. Consequently, we dynamically emphasize on optimizing these critical domains. With the reduction in the largest gap, the overall data distribution gap diminishes, augmenting the model’s robustness across diverse data distributions. We constructed GRNs for 88 mouse and 68 human bulk data sources from various tissues and developmental stages, as well as cell-type-specific GRNs for 40 mouse and 14 human single-cell sources (Table S1). Specifically, to achieve the transfer of GRNs across different tissue types, we treated each tissue in the GRN database as a domain. We calculated the distance between any two domains in the feature space, and selected a top percentage of the domains with the furthest distances as critical domains (these domains have large domain shifts that impedes the learning of domain-invariant knowledge) for generalization (Fig 2C and Methods). After identifying the critical domains, we used the Mixup strategy for data augmentation to reduce the distance between the critical domains (Fig 2D-E). After multiple rounds of dynamic selection of critical domains, data augmentation, and generalization, the model finally converged (Fig 2F). The same CDO strategy can also be applied to generalize GRN transfer learning across different developmental stages, cell types, and species.

### Performance of GRN Transfer Generalization Model

To evaluate the performance of our generalized model, we trained it using the single-cell-derived GRNs in our generated database and compared its performance with three state-of-the-art domain generalization methods: Domain-Adversarial Neural Networks (DANN), Mixup and CDO-MMD (Maximum Mean Discrepancy with CDO) (Ganin et al., 2016; Gretton et al., 2012; Xu et al., 2020). DANN introduced an adversarial training objective where it used a min-max optimization to learn invariant knowledge across domains. However, this method treated all domains equally, it may neglect the worst-case that is more crucial for model robustness. Random Mixup also has similar issue. CDO-MMD adopts Maximum Mean Discrepancy (MMD) instead of Mixup between critical domains since MMD is a widely used metric for adapting two domains. This comparison aims to evaluate the compatibility of our approach. The results showed that our model achieved a network correlation of nearly 95% (R^2^, see Methods) when compared with the true GRNs in the transfer of GRNs across different cell types in human or mouse single-cell data (Fig 3A). Various performance indicators, such as the root mean square error (RMSE), the mean absolute percentage error (MAPE), the area under the ROC curve (AUROC), were slightly better or comparable to the other three models. We also compared the predicted top 10,000 regulatory relationships by our model for a specific single-cell population with the top 10,000 edges generated by PECA2. The comparison showed that approximately 70-75% of the regulatory relationships were consistent between the two methods (Fig 3B). Overall, our results demonstrate the effectiveness and generalization capability of our approach for constructing cell-type-specific GRNs using transcriptomic data alone.

**Figure 3.**
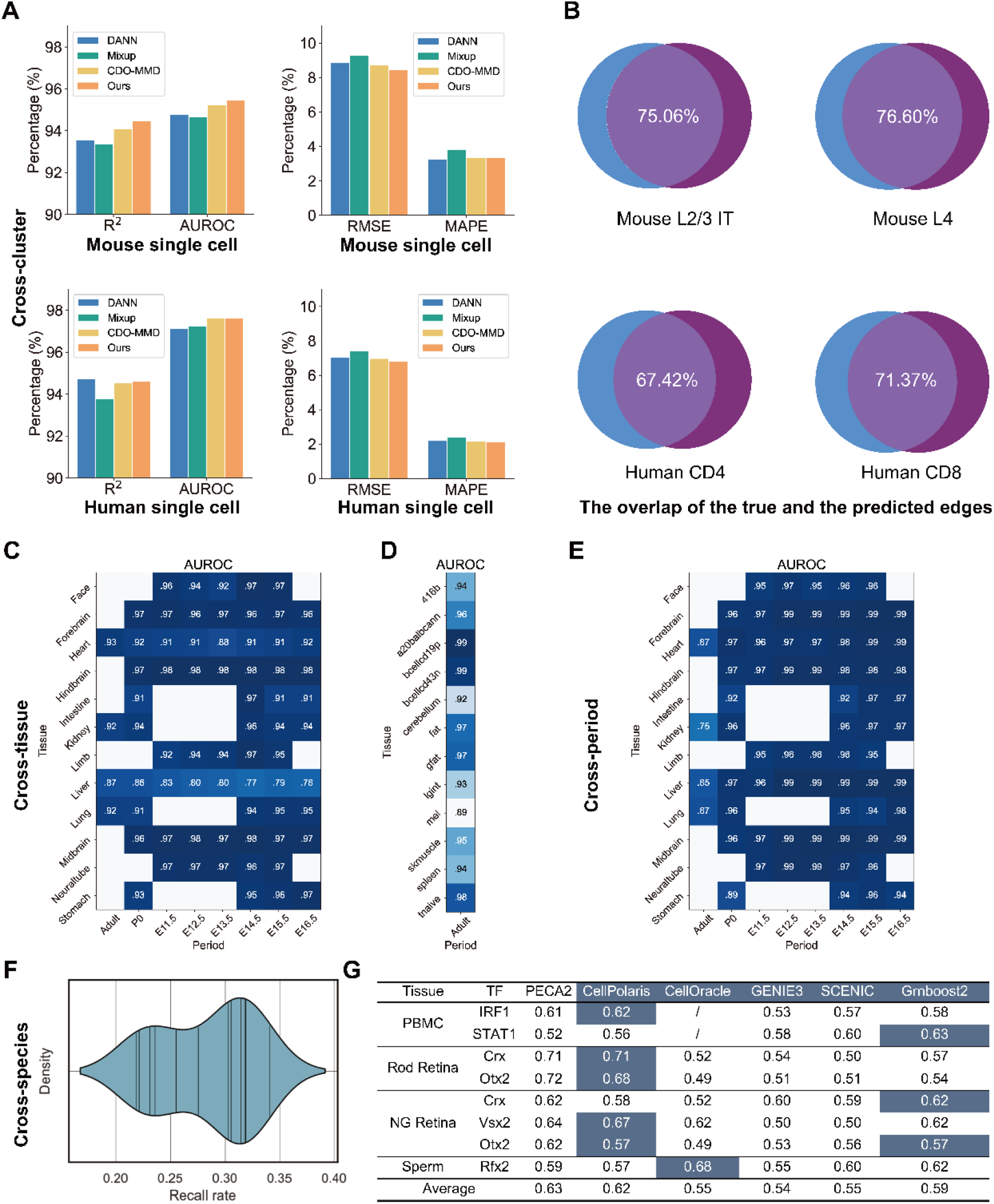
Performance of CellPolaris. **(A)** Cross-cluster evaluation on human and mouse single-cell RNA-seq data: Our model is trained using single-cell-derived GRN from PECA2 and compared with three state-of-the-art domain generalization methods on both human and mouse single-cell RNA-seq data. Each sc cluster is treated as a distinct domain, leaving one cluster out as the target domain. The model is then trained on the remaining source tissues and directly tested on the target to evaluate its generalization performance. **(B)** Comparison of GRN edges between PECA2 and model predictions: We conduct a thorough comparison between our model’s predictions and PECA2’s results for 4 human and mouse cells. The terms L2/3, IT (Intratelencephalic) neurons, and L4 are commonly used in neuroscience to classify specific layers and types of neurons in the mammalian brain; CD4 and CD8 are helper T cells and cytotoxic T cells, respectively. We evaluate the top 10,000 regulatory relationships predicted by our model and compare them with the top 10,000 edges generated by PECA2. **(C-D)** Cross-tissue evaluation: Similar to the cross-cluster evaluation, we treat tissues as separate domains. The model is trained on several source tissues and evaluated on data from the leave-one-out tissue to assess its ability to generalize across tissues. **(E)** Cross-period evaluation: Similar to the cross-tissue evaluation, we treat different developmental periods as separate domains. The model is trained on several source developmental periods and evaluated on data from the leave-one-out period to assess its ability to generalize across periods. **(F)** Cross-species evaluation: This evaluation poses a more challenging task, as it involves treating data from different species as distinct domains. We train the model using mouse single-cell RNA-seq data and perform inference on human single-cell RNA-seq data with homologous genes. **(G)** Comparison results on the ChIP-Atlas dataset: We compare the performance of our model with several state-of-the-art methods, including PECA2, CellOracle, Genie3, SCENIC, and GRNboost2, using the Chip-Atlas dataset. This analysis provides valuable insights into the effectiveness of our approach compared to other existing techniques.

Subsequently, we evaluated the accuracy of our model in predicting GRNs across different tissues and developmental stages using bulk data from mouse. During model training, we treated each tissue as a domain and used a leave-one-out setting, leaving one tissue as the target domain for testing and using other non-similar tissues as the source domain for training. We trained a model on the source domain and directly tested it on the target domain to evaluate the effectiveness of cross-tissue capability. The results showed that our model achieved a general accuracy of over 90% (AUROC) in predicting GRNs across different tissues (Fig 3C-D) and over 95% in predicting GRNs across different developmental stages (Fig 3E). In addition, we attempted to construct a direct transfer learning model for cross-species prediction without using any regulatory relationship information from the target species. This was to address the issue of the lack of high-confidence GRNs for training when predicting GRNs for non-model species. We used GRNs derived from all 30 single-cell sources in mouse to predict human single-cell data, and approximately 30% of the regulatory relationships among the top 10,000 homologous genes were accurately predicted (recall rates), indicating the potential usefulness of our model in cross-species prediction (Fig 3F). However, further optimization is still required.

Finally, we evaluated the performance of our model-generated GRNs by comparing them with those generated by existing GRN inference methods using ChIP-Seq data from ChIP-Atlas (Zou et al., 2022) as a benchmark (see Methods). We systematically compared the target genes of eight transcription factors across four cell types and found that CellPolaris achieved similar accuracy metrics (AUROC) to PECA2, which requires both ATAC-Seq and RNA-Seq data, and outperformed other reported methods (Fig 3G) (Aibar et al., 2017; Duren et al., 2020; Huynh-Thu et al., 2010; Kamimoto et al., 2023).

Our results indicate that CellPolaris is a reliable and accurate approach for predicting cell-type-specific GRNs using only transcriptomic data and can serve as a valuable tool for studying gene regulation in different cell types and tissues.

### Prediction of master TFs during cell fate transitions

The role of transcription factors (TFs) in cell fate transitions is well-established. Analyzing the regulation of target genes by these TFs is crucial for cell fate reprogramming beyond the simple selection of highly expressed or specific transcription factors. Drawing on this hypothesis, we utilized our established generalized transfer learning model to construct cell type-specific GRNs for the source and target cells in the reprogramming process. We identified target cell-specific differential GRNs by subtracting GRNs using established methods. We ranked all differentially expressed TFs from differentially expressed GRNs based on their expression levels in the source and target cells, the number of downstream target genes, and the proportion of downstream target genes that are differentially expressed between the two cell types (Fig 4A). We validated our predictions in three previously reported reprogramming or transdifferentiation systems (Graf, 2011; Rackham et al., 2016). Our results demonstrated that the majority of the top-ranked TFs were included in the five reported combinations of reprogramming factors (Fig 4B). However, some factors were ranked lower, as they may only accelerate the reprogramming process rather than being essential for cell fate transition (Nakagawa et al., 2008). We analyzed the top ten ranked TFs in each cell fate transition system and found that most of them have been reported to enhance reprogramming efficiency or be used for reprogramming with different combinations. Even for the TFs that have not been previously reported, they generally belong to important gene families, suggesting functional compensation (Fig 4C-D). Lastly, we showed that many of the top-ranked transcription factors regulate more than 10% of differentially expressed genes, indicating their importance in the reprogramming process (Fig 4E-G).

**Figure 4.**
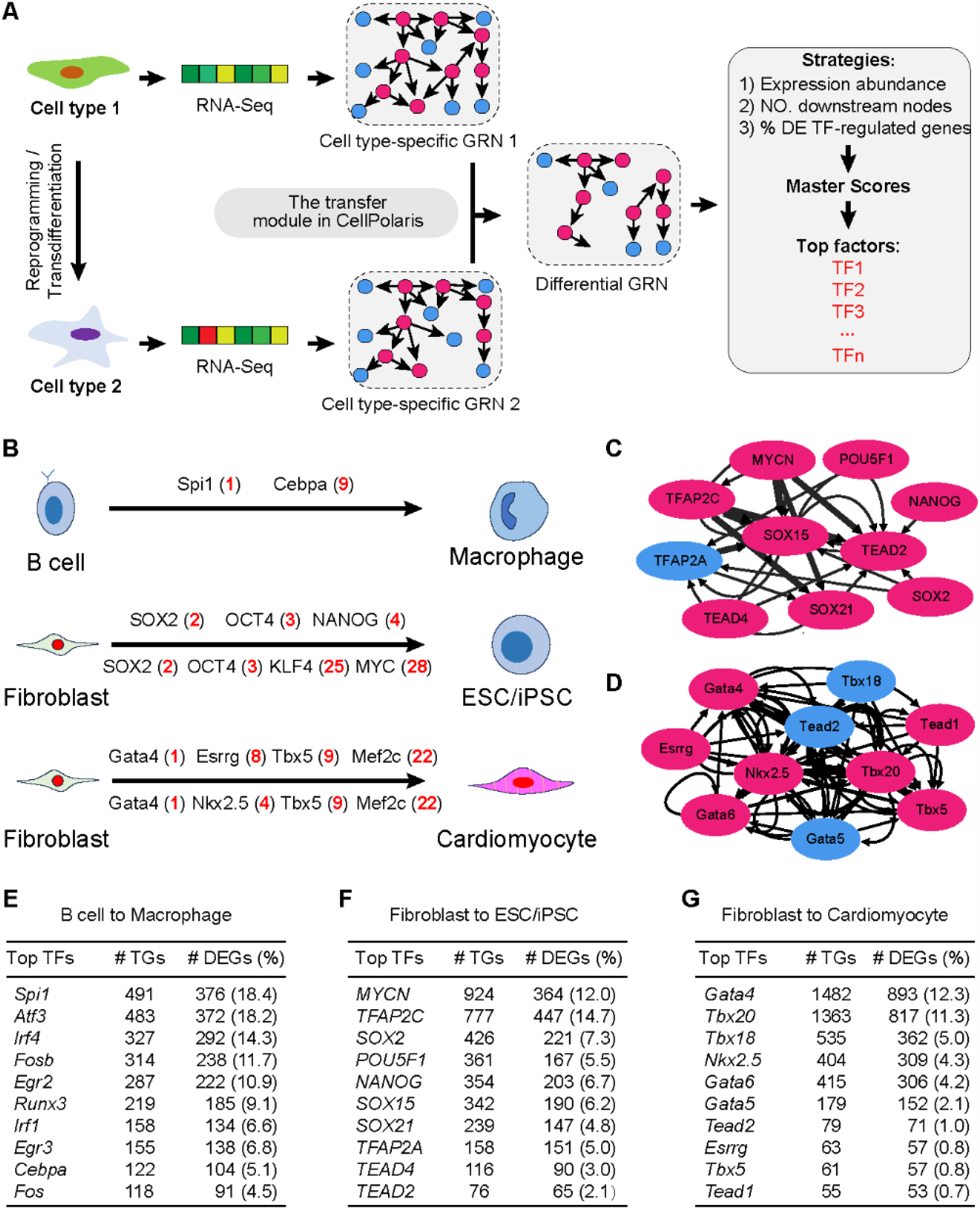
Prediction of master transcription factors in cell fate transitions. **(A)** Flowchart of master transcription factor (TF) identification and ranking. The GRN transfer generalization model was used to transform source and target cell transcriptome data into cell-type-specific GRNs. Differential TFs were identified by subtracting GRNs between cell types and scored based on expression abundance, downstream node numbers, and the number of differentially covered genes in the downstream nodes. **(B)** The model predicts the ranking of TF combinations known to be involved in reprogramming or transdifferentiation systems. The predicted rank of each factor is shown in parentheses. **(C)** The top 10 predicted TFs for MEF reprogramming into iPSCs by the model are shown, with red highlighting indicating TFs that have been used in fate conversion or validated to enhance reprogramming efficiency. **(D)** The top 10 predicted TFs for MEF transdifferentiation into cardiomyocytes by the model are shown, with red highlighting indicating TFs that have been used in fate conversion or validated to enhance reprogramming efficiency. **(E-G)** Statistical information on the model’s prediction of the top 10 master TFs regulating target genes in each cell fate transition system, as well as the percentage of differentially expressed target genes between the two cell states out of all differentially expressed genes.

Overall, we have demonstrated the applicability of our generalized transfer learning model in predicting cell fate transition factors with high accuracy, which can be applied to yet-unrealized cell fate transitions.

### Simulating gene perturbations during the process of round spermatid differentiation

In addition to predicting cell fate transition factors, GRNs have another important use for predicting the effects of gene perturbations during differentiation or reprogramming processes. Furthermore, high-resolution simulations of the effects of perturbations of important transcription factors on the differentiation process can be conducted along a single-cell developmental trajectory. To this end, we constructed a probabilistic graphical model to simulate TF perturbations at the single-cell level. Firstly, scRNA-Seq data was subjected to cluster analysis, and the cells in each cluster were transformed into pseudo-bulk data for generalized transfer of cluster-specific GRNs. The resulting GRNs were used to construct a probabilistic graphical model, which used the distribution parameters estimated from the cluster-specific nodes to infer the maximum probability distribution for all nodes. Based on this model, we predicted downstream target gene perturbations by setting the TF to zero. Finally, we adopted a similar strategy to the published CellOracle work and confirmed the effect of TF perturbations on cell developmental processes by calculating the consistency between simulated knockdown perturbations and direction of cell differentiation (Fig 5A).

**Figure 5.**
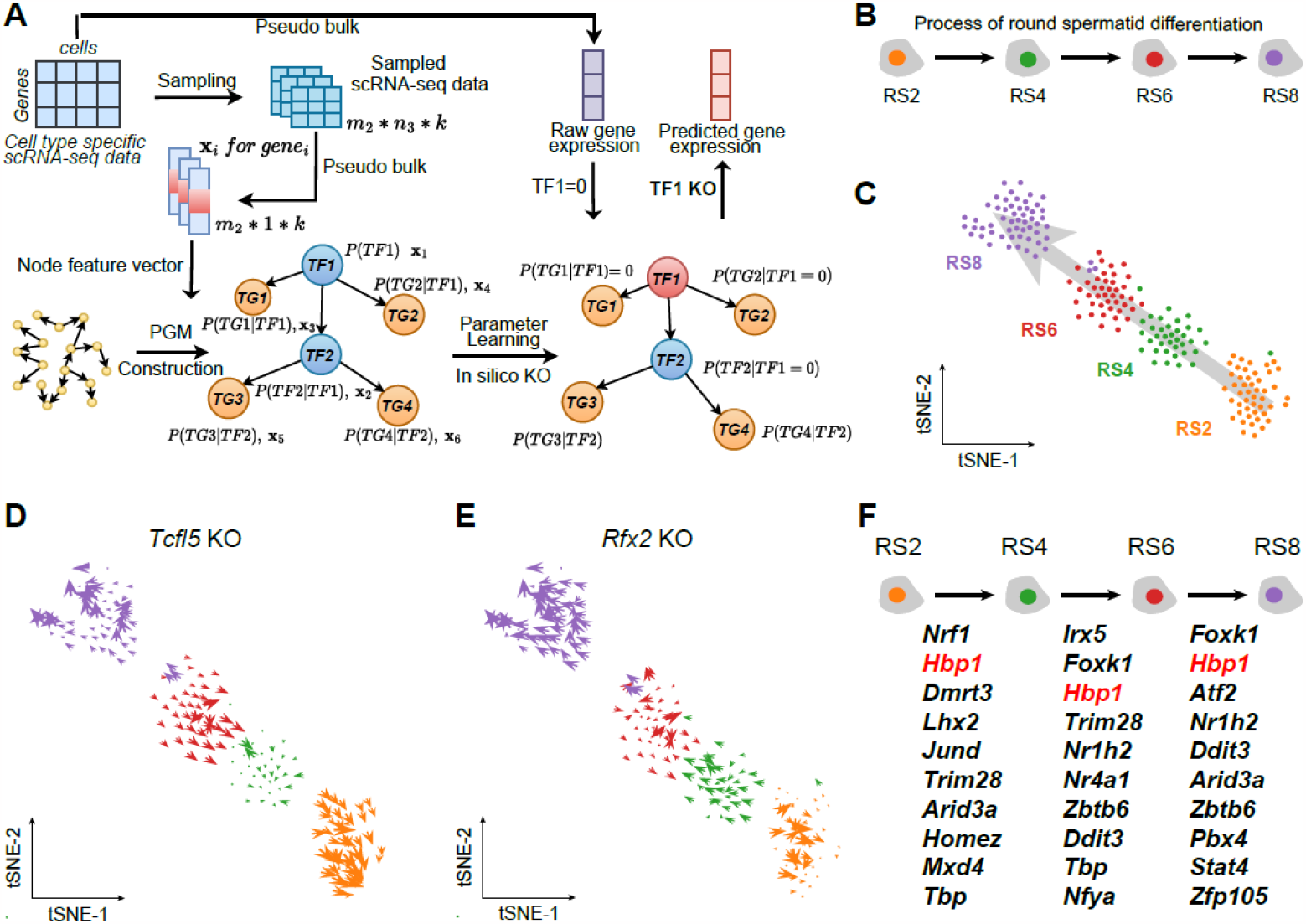
Simulation of the impact of transcription factor perturbation on cell differentiation progression. **(A)** Flowchart of the construction process for the probability graphical model (PGM) used to simulate transcription factor perturbation. Single-cell expression matrix was sampled to estimate the expression value distribution of each gene in the same cell cluster. The expression matrix of the same cell cluster was converted to pseudo-bulk data and used to predict the GRN with regulatory weights through the generalized transfer model. The PGM of gene expression was constructed by combining GRN and gene expression values, and parameter learning was performed. The downstream target gene expression changes were inferred through the model by setting the TF expression value to zero. **(B)** Diagram illustrating the process of round spermatid differentiation. RS2 represents steps 1-2 spermatids, RS4 represents steps 3-4 spermatids, RS6 represents steps 5-6 spermatids, and RS8 represents steps 7-8 spermatids. **(C)** t-SNE visualization of scRNA-Seq data from different stages of round spermatid differentiation, with arrows indicating the direction of differentiation. **(D)** Simulation of the impact of *Tcfl5* gene knockout on the development of round spermatids. Arrows pointing in the direction of differentiation indicate promotion, while arrows in the opposite direction indicate inhibition. **(E)** Simulation of the impact of *Rfx2* gene knockout on the development of round spermatids. Arrows pointing in the direction of differentiation indicate promotion, while arrows in the opposite direction indicate inhibition. **(F)** List of top 10 TFs that inhibit the differentiation process in round spermatids after gene knockout simulation at each stage. Factors that are present in all three stages are highlighted in red.

We applied the model to analyze single-cell data from the process of round spermatid differentiation after meiosis (Fig 5B) (Chen et al., 2018). By re-analyzing the single-cell transcriptome data from round spermatid differentiation, we constructed its differentiation trajectory (Fig 5C). Firstly, we measured the effects of two transcription factors that had been reported to regulate the process of round spermatid differentiation. The results suggested that knocking out both of transcription factors would reverse the differentiation trajectory of round spermatids, inhibiting differentiation (Fig 5D). Our prediction results were consistent with the results of CellOracle’s linear regression model. Sequentially, we identified important regulatory TFs involved in the differentiation of round spermatids, some of which were important in all three stages (Fig 5E-F). It is worth noting that the absence of many of these genes would result in early developmental arrest in animal models. Hence, our model holds great significance in guiding the study of gene functions in late-stage development processes.

In summary, we have established a probabilistic graphical model for simulating transcription factor perturbations during cell differentiation processes. We applied this model to simulate transcription factor perturbations during round spermatid differentiation and found that the results were consistent with those of genetic knockout animal models.

## Discussion

Constructing gene regulatory networks (GRNs) is critical for understanding the mechanisms of transcription factor interactions and organismal development. Here, we present CellPolaris, a generalized transfer learning model that generates state-specific gene regulatory networks (GRNs) solely from RNA-Seq data, leveraging existing high-confidence GRNs derived from both ATAC-Seq and RNA-Seq data. Compared to existing transcriptome-only-based software, such as GENIE3 and Grnboost2 (Aibar et al., 2017; Huynh-Thu et al., 2010), CellPolaris reasonably demonstrates significant performance gains. Furthermore, compared to other tools, such as PECA2, SCENIC, CellOracle, and DeepTFni, CellPolaris does not rely on any additional information beyond transcriptomics data, and its performance is not significantly reduced but rather improved (Aibar et al., 2017; Duren et al., 2020; Kamimoto et al., 2023; Li et al., 2022). One essential advantage of our proposed generalized transfer learning framework is its excellent scalability, and all high-confidence GRNs generated by existing GRN generation software can be used to improve CellPolaris’s performance.

As a deep learning framework, our model allows the use of embedding information generated by other models as prior knowledge. Several foundation models based on single-cell data have been developed, such as scGPT, Geneformer, and scFoundation (Haotian et al., 2023; Minsheng et al., 2023; Theodoris et al., 2023). These models utilize transformer architecture, pre-trained on millions of single-cell data, and generate embedding information for downstream tasks. The pre-trained information from these foundation models can be utilized to build and optimize transfer networks in CellPolaris. Moreover, the gene regulatory networks (GRNs) generated by CellPolaris can serve as prior biological knowledge for the pre-training process of these large models. This integration of prior knowledge can enhance the accuracy and efficiency of the pre-training process, ultimately leading to better downstream analysis and interpretation.

Regarding the downstream tasks involved in the gene regulatory networks (GRNs) produced by CellPolaris, we focus on two aspects: cell fate transition master factors and simulating the effects of TF perturbations on developmental processes. In terms of master factor prediction, our model utilizes a simple ranking scoring strategy, which achieves good performance compared to existing core factor prediction tools. Our strategy considers both the coverage of differentially expressed genes and the number of downstream target genes of TFs, demonstrating excellent scalability. For the perturbation prediction task based on probabilistic graphical models, our approach, compared to CellOracle, does not rely on the regulatory network derived from ATAC-Seq data and preserves the regulatory weights of TFs on downstream target genes, providing a higher level of model interpretability. It is worth noting that the downstream tasks of CellPolaris are expandable. For example, the GRN information generated by our model can be used for single-cell type identification or multi-modal omics data integration.

Finally, we acknowledge that our proposed generalized transfer learning framework has not demonstrated significant performance improvements at the cross-generational level compared to existing GRN construction methods. We suspect that this may be due to the limited availability of high-confidence GRNs used as training data in our study. In conclusion, we have successfully developed CellPolaris, a generalized transfer learning model, to generate cell type-specific GRNs solely relying on transcriptomic data. The model may provide insights into the mechanisms governing cell fate regulation and cell reprogramming, paving the way for further advancements in the field of gene regulatory network analysis

## Methods

Detailed methods are provided as separate chapters in the supplementary material of this paper, which include the following:

scRNA-seq data preprocessing.

Generate cluster pseudo bulk RNA-seq data from single-cell data.

PECA2 GRN construction.

Generalized GRN construction via critical domain optimization.

Ground truth network for GRN benchmark.

PGM-GRN algorithm.

*In silico* KO prediction method.

Differential Network Analysis and Visualization.

## Data and code availability

All primary data presented in this study have been deposited in a public database, and all codes have been uploaded to GitHub. After the official publication of the article, the data will be made publicly available, or researchers can directly contact the authors for access.

## Acknowledgements

We would like to thank Dr. Baoyang Hu and Dr. Wei Li for Their invaluable comments and guidance. We thank Chuanyang Zhang, Jiajia Wang, Chengrui Wang, Yan Chen, Yang Wang, Qirui Gu, Shan Zong, Huimin He from Computer Network Information Center, Chinese Academy of Sciences for their work in building website interface for model applications and model visualization. We also appreciate Dr. Wei Yang, Xuewei Yuan and Jie Zhang for their outstanding management support. We thank Beijing Institute for Stem Cell and Regenerative Medicine for the support provided. This work was also supported by CAS Project for Young Scientists in Basic Research, Grant No.YSBR-076.

## Author contributions

Xin Li, Yong Wang, Yiqiang Chen, Shihua Zhang, Yuanchun Zhou, Ming-Han Tong, Xuezhi Wang, Hongmei Wang and Ge Yang supervised the project. Xin Li, Yong Wang, Yiqiang Chen, Shihua Zhang, Yuanchun Zhou, Ming-Han Tong and Guihai Feng conceived and designed the study. Guihai Feng, Xin Qin, Jiahao Zhang, Wuliang Huang, Yiyang Zhang, Wentao Cui, Yao Chen and Shirui Li designed the algorithm and performed the experiments. Wenhao Liu, Yao Tian, Yana Liu, Jingxi Dong and Ping Xu were involved in analyzing sequencing data. Zhenpeng Man, Guole Liu, Zhongming Liang, Xiaodong Yang, Xinlong Jiang and Pengfei Wang provided assistance in designing the model. Guihai Feng, Wuliang Huang, Jiahao Zhang, Yiyang Zhang, Wentao Cui, Xin Qin and Shirui Li wrote the manuscript. The X-COMPASS Project Consortium members collaborated in the paper discussions. All authors read and approved the final version of the manuscript.

## APPENDIX

### The X-Compass Consortium Members

**Institute of Zoology, Chinese Academy of Sciences**

Xin Li, Hongmei Wang, Baoyang Hu, Wei Li, Fei Gao, Jingtao Guo, Leqian Yu, Qi Gu, Weiwei Zhai, Zhengting Zou, Guihai Feng, Wenhao Liu, Yao Tian, Chen Fang, Jingxi Dong, Yana Liu, Jingqi Yu, Wenhui Wu, Xinxin Lin, Cong Li, Yu Zou, Yongshun Ren, Fan Li, Yixiao Zhao, Yike Xin, Longfei Han, Shuyang Jiang, Kai Ma, Qicheng Chen, Haoyuan Wang, Huanhuan Wu, Chaofan He, Yilong Hu, Shuyu Guo, Yiyun Li

**Computer Network Information Center, Chinese Academy of Sciences**

Yuanchun Zhou, Yangang Wang, Xuezhi Wang, Pengfei Wang, Fei Li, Zhen Meng, Zheng Li, Zaitian Wang, Ping Xu, Wentao Cui, Zhilong Hu, Huimin He, Shan Zong, Jiajia Wang, Yan Chen, Chunyang Zhang, Chengrui Wang, Qingqing Long, Ran Zhang, Meng Xiao, Qinmeng Yang, Zijian Wang, Yining Wang

**Institute of Computing Technology, Chinese Academy of Sciences**

Yiqiang Chen, Yi Zhao, Xiaodong Yang, Dechao Bu, Xin Qin, Jiaxin Qin, Chaohui Yang, Chenhao Li, Zhufeng Xu, Zeyuan Zhang, Xiaoning Qi, Shubai Chen, Wuliang Huang, Boyang Zhang, Yaning Li

**Institute of Automation, Chinese Acadery of Sciences**

Ge Yang, Jing Liu, Guole Liu, Jie Jiang, Xingjian He, Liqun Zhong, Yaoru Luo, Jiaheng Zhou, Zichen Wang, Ziwen Liu, Ao Li, Teng Wang, Yiming Huang, Handong Li

**Academy of Mathematics and Systems Science, Chinese Academy of Sciences**

Yong Wang, Shihua Zhang, Jiahao Zhang, Yiyang Zhang, Shirui Li, Zhongming Liang, Zhenpeng Man, Kangning Dong

## Figures & Figure legends

**Figure S1.**
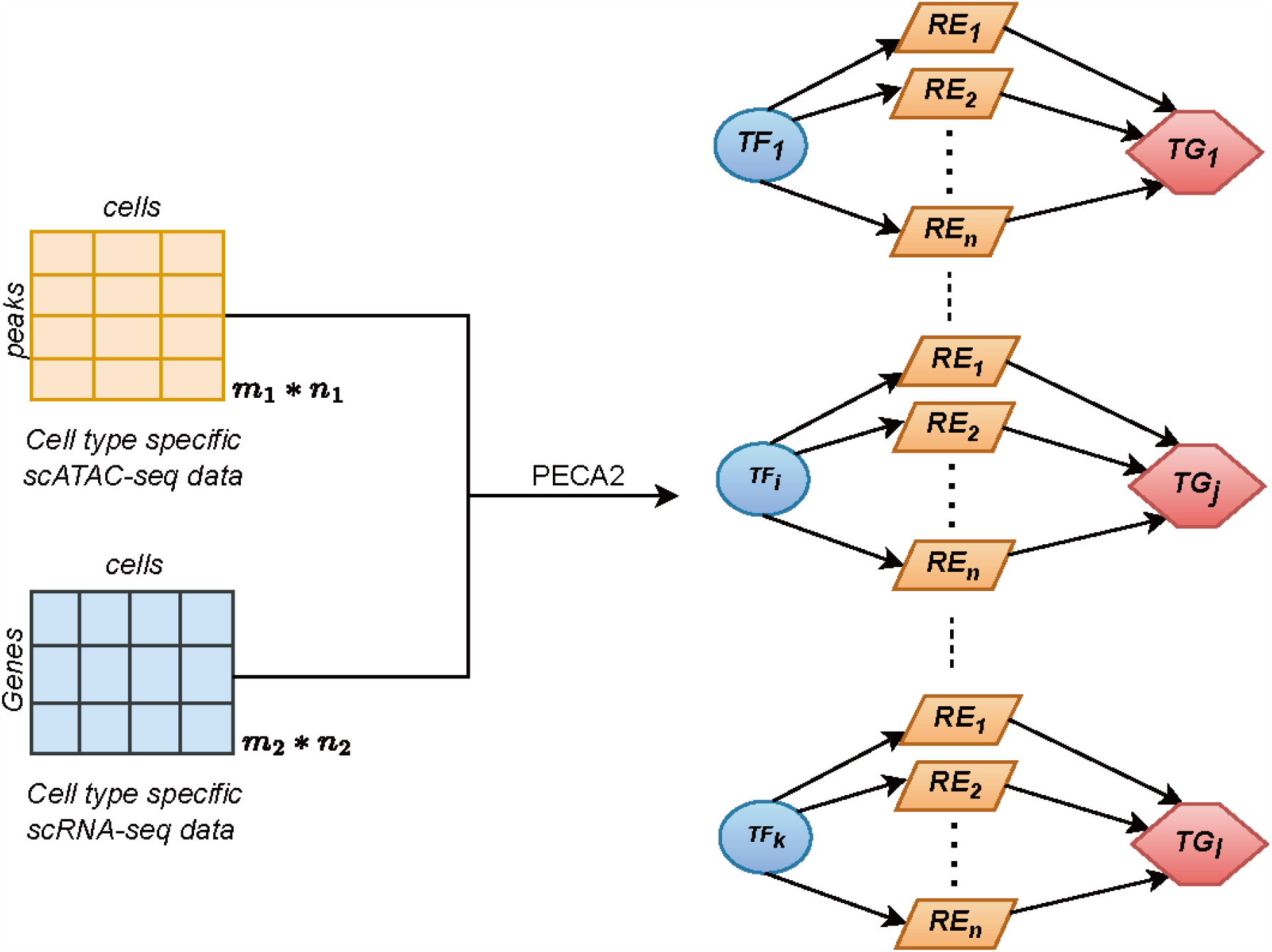
PECA2 generates GRNs with regulatory weights using RNA-Seq and corresponding state-specific ATAC-Seq data.

**Table S1. GRN tissue and developmental stage information used for training the CellPolis model**.

## Method

### scRNA-seq data preprocessing

For a count matrix of single-cell data, we usually use Scanpy^1^ for preprocessing. Firstly, we filter out cells with gene expression below 200 and cells with mitochondrial gene percentage exceeding 15%. Then, we linearly scale each cell based on the total UMI count to 10,000 by using scanpy.pp.normalize_total(adata) and then applying a log1p transformation to the expression values using scanpy.pp.log1p(adata), we detect 2000 highly variable genes using scanpy.pp.highly_variable_genes(adata, ntop_genes=2000). We further perform dimensionality reduction and clustering on the expression matrix for subsequent downstream analysis.

### Generate cluster pseudo bulk RNA-seq data from single-cell data

We first Segmenting the single-cell expression matrix into multiple sub-matrices based on cell types through clustering. We further process each sub-matrix for individual cell types. First, we calculate the total expression of each gene within a cell by summation. Then we can obtain the Fragments Per Kilobase Million (FPKM) values for the generated pseudo-bulk data as follows:

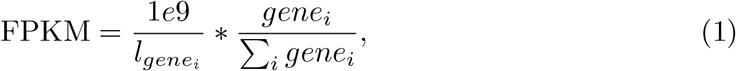

where *gene*_*i*_ is the summation of gene expression count values across all cells and 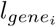 is the length of *gene*_*i*_.

### PECA2 GRN construction

The original PECA2 uses paired expression(RNA-seq) and chromatin accessibility data (ATAC-seq) as input to generate gene regulatory network by selecting active REs, specifically expressed TFs and expressed TGs in the context. In our scenario. We need to obtain a more refined gene regulatory network for each stage of cell differentiation, so we select well-labeled scRNA-seq data and well-labeled scATAC-seq data to make cell type specific PECA2 network. While the data is not well-labeled, we can run the Seurat CCA process to integrate scRNA-seq and scATAC-seq data. For each cell type, we sum scRNA-seq data and then calculate the FPKM as pseudo bulk RNA-seq data, and normalize the sum of scATAC-seq data as cell type openness data(2 normalization Methods, *counts of peak*_*i*_*/counts*_*mean*_, or *counts of peak*_*i*_*/background counts of peak*_*i*_. For ease of use, we select the former.) After this process, we get cell type paired psuedo bulk RNA-seq data and psuedo bulk ATAC-seq data to construct PECA2 network as follows:

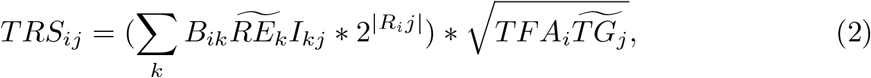

where *B*_*i*_*k* is motif binding strength of *TF*_*i*_ on *RE*_*k*_, which is defined as the maximal binding strength calculated by HOMER software of all of the binding sites on 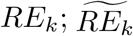 represent the accessibility of *RE*_*k*_; *I*_*k*_*j* represents the interaction strength between *RE*_*k*_ and *TG*_*j*_, which is learned from the PECA model on diverse cellular contexts; *R*_*i*_*j* is the expression correlation of *TF*_*i*_ and *TG*_*j*_ across diverse cellular contexts; *TFA*_*i*_ represents the activity of *TF*_*i*_ (geometric mean of normalized expression and motif enrichment score on open region); 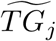 represents the normalized expression level of 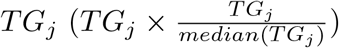.

We use the PECA2 edges and TRS as part of the input to construct the PGM model.

### Generalized GRN construction via critical domain optimization (CDO)

In the followings sections, we propose to construct cell-type-specific GRN by CDO.

### GRN generator

To predict the regulatory relationship between Transcription Factors (TFs) and Target Genes (TGs), first, we propose a Gene Regulatory Network(GRN) generator to build GRN with gene names and gene expressions. Inspired by Neural Collaborative Filtering (NCF) [5] that leverages a multi-layer perception to model the user–item interaction function, our GRN generator explores the interactions between TFs and TGs, as well as the interactions between their corresponding gene names, to generate the GRN. The key idea is to leverage the non-linear relationships learned by neural networks to capture the complex interactions between TFs and TGs. The GRN generator represents TF expression value *x*_*tf*_ and TG expression value *x*_*tg*_ as dense vectors through embedding layers, which capture the latent features of TFs and TGs. These embeddings are then fed into neural networks to learn the interaction patterns and make predictions. The prediction function can be formulated as:

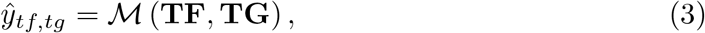

where *ŷ*_*tf,tg*_ is the predicted score for TF and TG, **TF** is the embedding vector for TF, and **TG** is the embedding vector for TG, ℳ is the whole model parameters.

Specifically, we utilize the gene IDs of TFs and TGs, denoted as *id*_*tf*_, *id*_*tg*_ respectively, as the input for both the Multi-Layer Perceptron (MLP) embedding layer *h*_*mlp*_ and the matrix factorization (MF) [4, 8] embedding layer *h*_*mf*_ to obtain the respective MLP and MF embeddings for the gene IDs, denoted as *h*_*mlp*_(*id*_*tf*_), *h*_*mlp*_(*id*_*tg*_) and *h*_*mf*_ (*id*_*tf*_), *h*_*mf*_ (*id*_*tg*_). Subsequently, we concatenate the MF embedding with the corresponding gene expression value as [*h*_*mf*_ (*id*_*tf*_) ∥**x**_*tf*_], [*h*_*mf*_ (*id*_*tg*_) ∥**x**_*tg*_] and then perform an element-wise product between the TFs and TGs. The resulting product is then fed into the MF-module to derive the MF features:

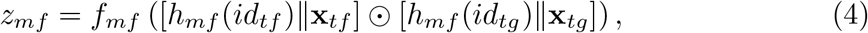

where *f*_*mf*_ (*·*) is a Multi-Layer Perceptron.

Then take **x**_*cat*_ = (**x**_*tf*_ ∥*h*_*mlp*_ (*id*_*tf*_) ∥(**x**_*tg*_ *−* **x**_*tf*_)∥(*h*_*mlp*_ (*id*_*tg*_) *− h*_*mlp*_ (*id*_*tf*_))) as the input of the MLP module to get MLP features *z*_*mlp*_:

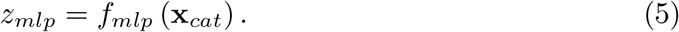

By concatenating the MF and MLP features as the input of regression layers, we can get the prediction.

### Improve domain diversity

Different genes are expressed to different degrees in different types of cells, and this results in the discrepancy of the overall gene expression distribution with various cell clusters or bulk types. In the following, we collectively refer to cell or bulk types as “different cell types”. Due to the specialized gene expression discrepancy in different cell types, we propose to rank the expression value and make this rank embedding as an input of our model to utilize the diversity of cell type, thus improving the generalization capability. To implement this, we rank all expression values in descending order for each PECA2 data of each cell type and obtain the corresponding gene indices in the list of all genes. Then, we judge whether the expression value is in the first quartile, the second quartile and the third quartile to help us understand the distribution and variability of the expression degree of genes in different cell types, and by comparing data points to the quartiles, we can assess their relative positions and distribution patterns within the overall dataset. Finally, we concatenate this ranking information with the aforementioned input for our model. Besides, we incorporate two types of domain features into the input based on the data type: whether it is from bulk, whether is from single cells. By adding these domain features, the domain-specific feature is enhanced and intro more diversity into the data space.

### Critical domain optimization

Regard each cell type data as a domain with each domain representing a point in the data space. The similarity between a pair of domains can be seen as the relationship between them. We regard domains as vertex and the similarities as edges among them, then we can build a domain graph that can draw the relationship of all cell types. The core of our method is to fill in the space between domains to smoothly enlarge the diversity of data space, especially between domains those are far from each other, i.e. the worst cases. These domains are critical for generalization. By improving the critical domains, the model focuses more on the knowledge that is hard to learn. The similarity is calculated every epoch, and the critical domains are changing dynamically. With the smooth bridging between less similar domains, the edges among all domains are constructed with more diversified mixup samples and the domain graph becomes more complete. By optimizing critical domains, the robustness and generalization capability of the model is enhanced.

We first calculate the distance (the larger the distance, the smaller the similarity) between training domain pairs by utilizing Maximum Mean Discrepancy (MMD) [3] which is a widely used distance metric in transfer learning. The distance between two domains 𝒟_*s*1_ and 𝒟_*s*2_ is:

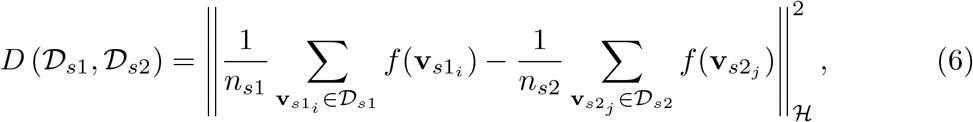

where *D* ((·, ·) is the distance between two domains. **v**_*s*_ and **v**_*t*_ is the input data from source domain and target domain, where **v** = [**x**_*tf*_, **x**_*tg*_, *id*_*tf*_, *id*_*tg*_]. *n*_*i*_ and *n*_*j*_ are cardinalities of these two domains. *f* (·) is the feature map which maps the original instances into the RKHS ℋ.

We rank the edge values of the domain graph in descending order. When the distance is in the threshold of top-*σ* ratio, we believe that the two domains are far apart that the gap between them need to be filled and rebuild the edge between them. To smoothly reduce the distribution discrepancy between two far domains, we propose to use an alternative data augmentation technique. Specifically, we utilize Mixup [10] as the augment technique to smoothly blur the boundaries of domains. Mixup is a simple but effective data augmentation method by making linear interpolations on random pairs of samples. Given two random samples (**x**_*i*_, *y*_*i*_) and (**x**_*j*_, *y*_*j*_), the mixup between them can be formulated as:

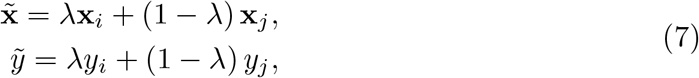

where *λ ∼ Beta* (*α, α*) and *α ∈* (0, *∞*) are hyperparameters.

In our generalized GRN construction problem, the original mixup can not be used directly. As we focus on learning the regulator weight from the 4 tuples of data [**x**_*tf*_, **x**_*tg*_, *id*_*tf*_, *id*_*tg*_], i.e., the expression value pairs of TFs and TGs and their corresponding gene IDs. Thus, we adopt mixup in the TFs and TGs and their embeddings with the regulator weights. For each *k* iterations, we get the top-*σ* ratio of critical domain pairs. Then random pick sample pairs 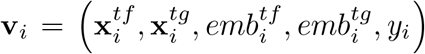 and 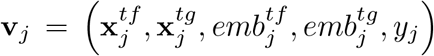 from two domains to mixup. This process can be formulated as follows:

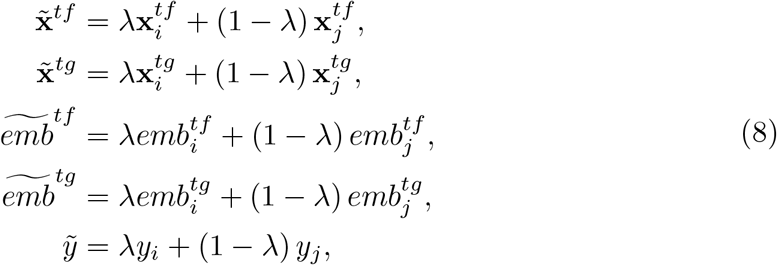

where *emb*^*\**^ is the embedding of [**x**^*tf*^, *idemb*^*tf*^, **x**_*tg*_*−***x**_*tf*_, *idemb*^*tg*^*−idemb*^*tf*^], and *idemb*^*\**^ is the embedding of gene ID.

Thus, the learning optimization is to minimize the regression loss of the original data and the loss of virtual data genrated by Mixup between the critical domain pairs as Equ. 9.

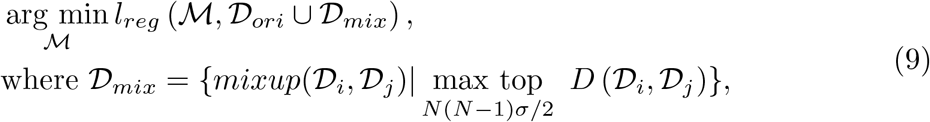

where the basic loss function *l*_*reg*_ is MSE loss and 𝒟_*ori*_ and 𝒟_*mix*_ are domains of the original and augmented data. *σ* is the ratio of the critical domains from all *N* source domains. *S* denotes the critical domain set which cardinality is *N* (*N−*1)*σ/*2.

By filling up the space between domains, the latent similarity between domains are enlarge and the graph are dynamically rebuild. Thus, the model can better learn the general features to achieve more robust and generalized inference on new data.

### Ground truth network for GRN benchmark

To verify whether the transfer learning network can accurately predict cell-type-specific gene regulatory networks, we conducted benchmark tests by comparing it with several known gene regulatory network inference methods, including PECA2 [2], CellOracle [7], GENIE3 [6], GRNboost2 [9], and SCENIC [1].

GENIE3 is a regression model based on Random Forests or ExtraTrees. GRN-Boost2 is a high-performance algorithm that utilizes gradient boosting and is based on the GENIE3 framework for inferring regulatory networks. The SCENIC algorithm combines a tree-based algorithm for inferring gene regulatory networks with information about transcription factor binding.

To generate the cell-type-specific ground truth gene regulatory networks, we utilized the ChIP-Atlas database^2^. For each specific cell type or tissue, we downloaded TF ChIP-seq data as bed files from the ChIP-Atlas datasets. We then selected TF peaks present in the ChIP-seq data. The corresponding TF-TG scores were downloaded from ChIP-Atlas and converted into a binary network, where each edge weight is either 1 or 0, indicating the presence or absence of ChIP-seq binding between genes. The conversion process involved the following steps: (i) selecting the top 100 genes based on the binding scores of MACS2 and STRING for each TF as positive edges in the binary network; (ii) deleting all the genes with non-zero binding scores from the target gene list for each TF. This process resulted in a gene regulatory network specific to the cell type or tissue of interest. For our benchmark tests, we selected eight regulatory factors across four specific cell types. The performance of network inference was evaluated using the area under the receiver operating characteristic (AUROC) metric.

### PGM-GRN algorithm

The **P**robabilistic **G**raphical **M**odel Based on **G**ene **R**egulatory **N**etworks algorithm consists of the following main steps: (1) Data processing and construct PECA2 network using paired bulk(or pseudo bulk) RNA-seq and ATAC-seq data from the same cell type.(2) Construct the graph whose nodes are genes and edges represent the regulatory relationships from PECA2 network.(3) Sample cells from scRNA-seq data for many times, and merge the cells into pseudo bulk RNA-seq for each time.(4) Each node gets its feature vector from (3), each edge gets its strength from PECA2 network.(5) Use the information in (4) to construct the Probabilistic Graphical Model and perform parameter training. (6) Do in silico TF knock out predictions and observe the direction of cell fate changes.

### PGM-GRN model overview

Probabilistic graphical model is a powerful tool for representing complex probability distributions in a compact and intuitive way. Bayesian network, is a probabilistic graphical model that represents a set of random variables and their dependencies. It consists of nodes, which represent variables, and directed edges, which represent causal relationships between the variables. Each node is associated with a probability distribution that describes the variable’s possible values given its parent variables. In our PECA2 network situation, we model genes as nodes and regulatory relationships as edges to get a directed graph construction. We need to make some assumptions and arguments to populate this graph.

#### Hypothetical description

In general, single-cell gene expression does not follow a normal distribution. We performed an operation to bring the gene expression distribution closer to the normal distribution. For one cell type, assume that there are *N* scRNA-seq cells, We randomly sample *N/*10 cells and merge them to pseudo bulk. The above process is repeated *S*(default: 50) times. The genes in the *S* samples should approximately follow a normal distribution. Next, we use the Central Limit Theorem to make some arguments for the hypothesis.

#### Lévy-Lindeberg Theorem

Let *{ξ*_*n*_*}* be an independent and identically distributed(i.i.d.) sequence of random variables such that E(*ξ*_*n*_) = *a < ∞*, 0 *<* Var(*ξ*_*n*_) *< ∞*, then *{ξ*_*n*_*}* satisfies the Central Limit Theorem, namely,

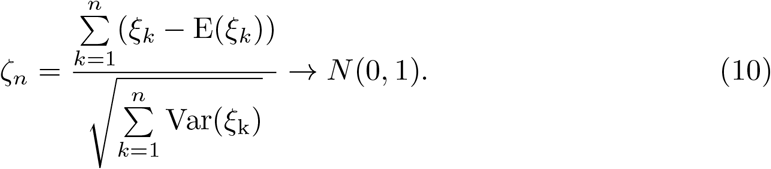

#### Hypothetical argumentation

For a single cell, the number of mRNA fragments detectable by the typical scRNA-seq method is much smaller than the number of mRNA fragments not detected. Therefore, when we sample cells and synthesize pseudo bulk, each gene fragment can be viewed as being obtained from a sufficiently large population. Assuming there are C sequencing fragments in our pseudo bulk data, for each fragment, the probability of it being *gene*_*i*_ is *p*_*i*_. Random variable defined as follows, *ξ*_*ki*_ *∼ B*(1, *p*_*i*_), *∀k ∈ {*1, 2, …, *C}*, E(*ξ*_*k*_) = *p*_*i*_, Var(*ξ*_k_) = *p*_*i*_(1 *− p*_*i*_). Brought into the theorem, we get

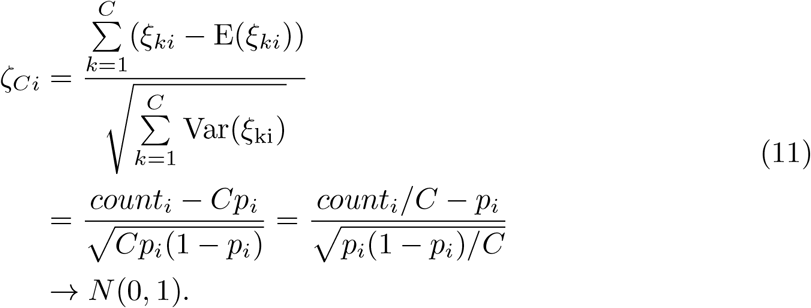

The FPKM for *gene*_*i*_ is proportional to *count*_*i*_*/C*, and our sampling policy in the same cell type reduce differences in counts between samples, so we can make an approximation assumption that the FPKM for 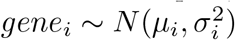.

#### PGM model objective function

According to the PECA2 network structure, we divide nodes into two categories: (1) top TFs, whose indegree are 0, recorded as *S*_1_; (2) others, consist of TGs and other TFs, recorded as *S*_2_. Our objective function is to learn the parameters that make observations the most probable, so as to obtain the conditional probability distribution between nodes, i.e.

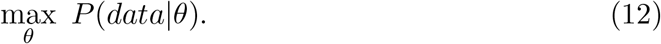

Distinguish between *S*_1_ and *S*_2_ through different probability forms, the objective function can be written as,

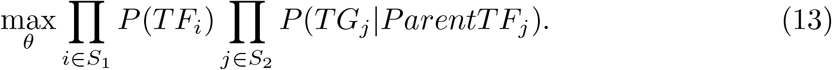

where, *TF*_*i*_ represents the FPKM value of *TF*_*i*_ in all samples; *TG*_*j*_ represents the FPKM value of *TG*_*j*_ in all samples; *ParentTF*_*j*_ represents the set of TFs who regulate *TG*_*j*_ from PECA2 network; the parameter set *θ* consists of each gene’s distributed parameter 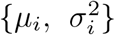 and TRS effect control parameter *{l*_*ij*_*}*. Next, we analogy the form of a binary Gaussian distribution to introduce how to bring in TRS and {*l*_*ij*_}.

#### Conditional distribution form of binary Gaussian distribution

Assume that **Y** *∼ N* (*μ*, Σ), Σ *>* 0(positive definite), where

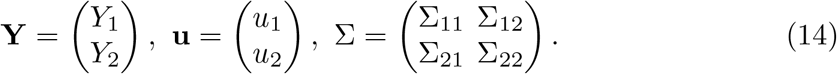

then given *Y*_2_ = *y*_2_, the conditional distribution of y1 is *N* (*μ*_1|2_, Σ_1|2_), its mean and variance are

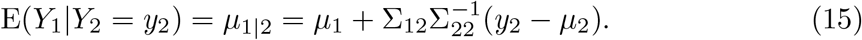

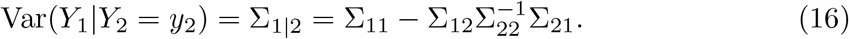

The correlation of gene expression (here, the covariance Σ_12_, Σ_21_) can not well indicate the regulatory intensity between TF and TG, and TRS is a more real regulatory intensity obtained according to the sequencing data and the inherent motif information of TF. Assume that 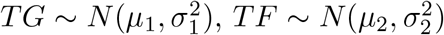, the product of TRS and *Cov*(*TF, TG*) is used instead of Σ_12_, Σ_21_, and *l* parameter is added to control the range of the regulatory intensity in the conditional distribution, i.e.

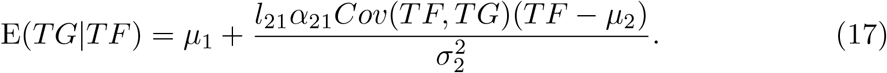

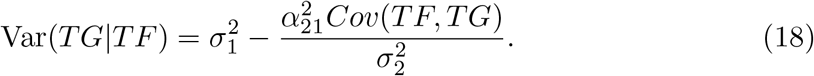

where, *α* corresponds to TRS, and default value range of parameter *l* is [−10, 10]. Simply look at the forms of formulas (4) and (5), the greater the regulatory intensity, the difference between TF expression and its mean value will lead to greater changes in TG expression. In addition, the greater the regulatory intensity, the smaller the variance of *TG*|*TF*, while the smaller the regulatory intensity, the variance of *TG*|*TF* tends to the prior of TG, which is consistent with biological common sense.

#### Objective function solving

After performing the above transformation, we obtain a conditional probability representation on the edge. Return to the conditional probability term of the objective function, we can expand it according to the Bayes formula. Assume that *ParentTF*_*j*_ = *{TF*_1_, *TF*_2_, …, *TF*_*n*_*}*, then

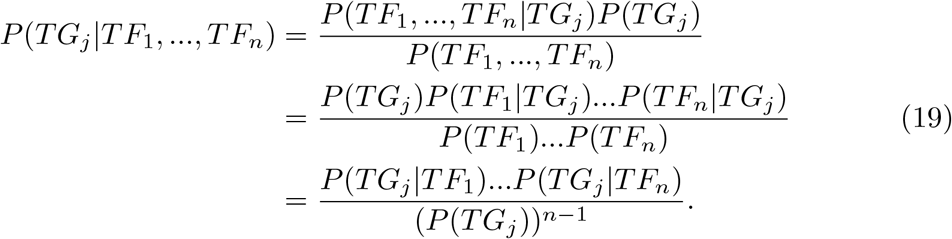

The second equal sign uses the assumption that *TFs* are approximately independent of each other, because the processes in gene regulation are mainly based on recognition after random collisions. After explicitly writing out each term in the objective function, we use stochastic gradient descent(SGD) to minimize the -log objective function, and the default number of iterations is 100.

### In silico KO prediction method

The purpose of constructing PGM and learning the parameters is to obtain the cell fate transfer and predict the expression value after TF knockout. Suppose we knock out *TF*_1_, its’ target gene *TG*_*j*_’s expression value can be estimated as follows:

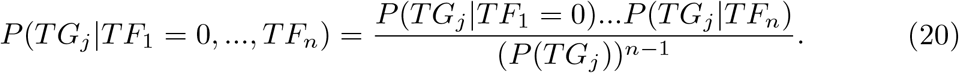

It is worth noting that in the hypothetical argumentation section, the variance of gene distribution is inversely proportional to the total number of gene fragments. Because the cells we sampled(default cell number: N/10) come from the same cell type(cell number: N), we can substitute the number of cells for the number of fragments. Therefore, under the default selection, we change the variance of gene distribution *σ*^2^ to *σ*^2^*/*10 for each gene in the KO task. In formula (6), we set *TF*_1_ to 0 and other *TFs* remain unchanged, so the only unknown is the value of TG to be predicted. Find the maximum value of equation (6), and its maximum point is the predicted value of *TG*_*j*_.

After traversing all the downstream *TG* of *TF*_1_, we obtained the first round predicted value of gene expression. *TF*_1_ and its downstream *TG* are changed and recorded as set *V*_1_. We can then make predictions about *TF*_1_’s second-order neighbors. For a TG, which is *TF*_1_’s second-order neighbor, use the changed value for *TF* in *V*_1_ and the original value for *TF* outside *V*_1_, and Find the maximum value of equation (6) to get the TG’s predicted value.

For cell fate transfer task, We calculated the expression value of each gene after knockout. The original expression vector is recorded as **e**_1_, the prediected expression vector is recorded as **e**_2_, and another cell type(c) expression vector is recorded as **e**_*c*_. Measured by cosine similarity, where will the cell state shift after knocking out a TF,i.e. cos *{***e**_2_*−***e**_1_, **e**_*c*_*−***e**_1_ *}*.

The higher the cosine value, the more likely the cell state is to transfer to cell type c after knocking out this TF.

### Simulate and draw the cell differentiation trajectory after knocking out a specific transcription factor

For simulation of cell identity, we developed our code by modifying Celloracle [7]. To simulate the differentiation direction of cells following perturbation, we first identify the cells belonging to each cluster based on the clustering results obtained previously. Then we calculate the FPKM expression value for each gene across all cells within a cluster. This can be done by taking the sum expression value of each gene within the cells of the cluster. Repeat the above step for each cluster, resulting in a pseudo-bulk expression matrix where each row represents a gene and each column represents a cluster. Use a probabilistic graphical model we can obtain the change in gene expression values, *deltaX*, within each cluster after knocking out a specific transcription factor (*TF*). After obtaining the *deltaX* for each cluster, we can assign the corresponding change value to each cell within that cluster. Then, we calculate the similarity between this change, *deltaX*, and the difference in expression vectors of the cell’s neighboring points minus its own expression vector. Please note that when calculating similarity, we only consider the target genes (*TG*) directly connected to the transcription factor (*TF*) in the gene regulatory network. The reason behind this approach is that when we considering the impact of knocking out a *TF* on changes in gene expression and subsequent cell fate changes, the direct influence is expected to be concentrated mainly on the *TGs* directly connected to the *TF* in the gene regulatory network. These *TGs* are likely to experience the most significant effects. Therefore, when calculating similarity, we only consider the effects on these *TGs*. The calculated similarity can then be converted into the probability of cluster-to-cluster transfer for the given cell and its neighboring cells. Multiplying this transition probability by the distance vector in the low-dimensional feature space between the cell and its neighboring cells, we can synthesize the final simulated differentiation direction of the cell.

### Calculate the cosine similarity to find the important TFs

To generate a perturbation score after knocking out a specific transcription factor, we can calculate the pseudotime gradient vector field and the inner product score. Here’s how it can be done: For cell fate transfer task, We calculated the expression value of each gene after knockout. The original expression FPKM vector of this cluster is recorded as **e**_1_, the *deltaX* expression vector is recorded as **e**_2_, and another cluster expression FPKM vector is recorded as **e**_*c*_. Then we can calculate the similarity between the change in (*deltaX*) within the cluster and the difference in FPKM values between adjacent clusters.

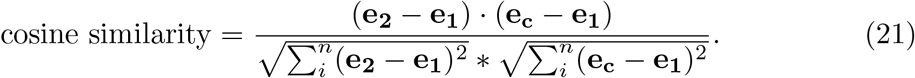

A positive value indicates that knocking out the transcription factor promotes the transition to the next cluster, while a negative value indicates an inhibitory effect.

### Differential Network Analysis and Visualization

Assuming there are two gene regulatory networks (PECA2 network or transfer learning network) *N*_1_ and *N*_2_, the specific network of *N*_2_ for *N*_1_ is calculated as follows.(1) Use the already generated PECA2 network (84 mouse network or 76 human network) as background. If a TF-TG regulatory relationship occurs more than 10 times, the edge is considered common and will be removed. (2) Select the top 5000 unique regulatory relationship in *N*_2_ ranked by the score as the differential network, delete the common edges in (1). (3) Calculate two ranks for TFs in the differential network. The first rank is the outdegree of the TF in the differential network, and the second one calculate a TF’s TG specific score as follows: *TG specific score for* 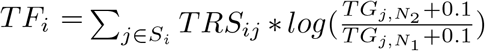. This specific score reflects the ability of a TF to regulate downstream differentially expressed genes. Rank this score in descending order as the second ranking. Add these two rankings and rank them again from top to bottom as the final ranking of TF in the differential network. (4) Select the regulatory relationship whose TF and TG are all in top 10 TF list to show the regulatory relationship between the top 10 TFs.

## Footnotes

1 https://scanpy.readthedocs.io/en/stable/

2 https://chip-atlas.org

