## Supplementary figures and images for "CellPolaris: Decoding Cell Fate through Generalization Transfer Learning of Gene Regulatory Networks"

### Fig S1

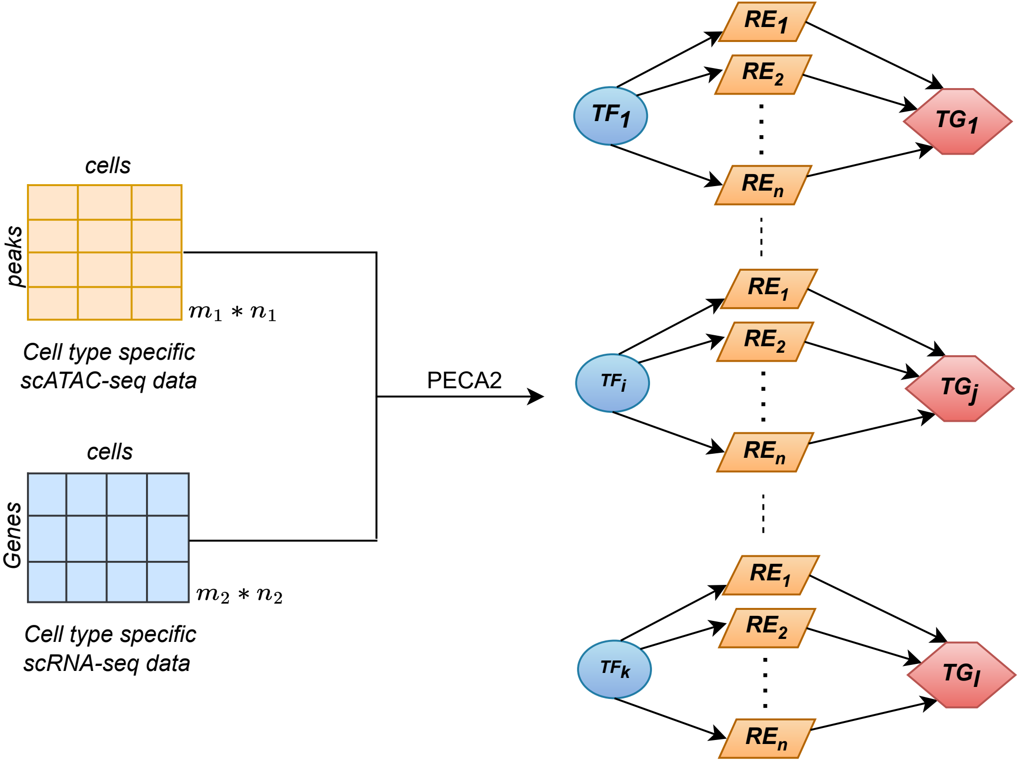
